# Dynamical state of the network determines the efficacy of single neuron properties in shaping the network activity

**DOI:** 10.1101/030700

**Authors:** Ajith Sahasranamam, Ioannis Vlachos, Ad Aertsen, Arvind Kumar

**Affiliations:** Bernstein Center Freiburg and Faculty of Biology, University of Freiburg, 79104 Freiburg, Germany.; Computational Biology, School of Computer Science and Communication, KTH Royal Institute of Technology, Stockholm, 10044, Sweden

## Abstract

Spike patterns are among the most common electrophysiological descriptors of neuron types. Surprisingly, it is not clear how the diversity in firing patterns of the neurons in a network affects its activity dynamics. Here, we introduce the state-dependent stochastic bursting neuron model allowing for a change in its firing patterns independent of changes in its input-output firing rate relationship. Using this model, we show that the effect of single neuron spiking on the network dynamics is contingent on the network activity state. While spike bursting can both generate and disrupt oscillations, these patterns are ineffective in large regions of the network state space in changing the network activity qualitatively. Finally, we show that when single-neuron properties are made dependent on the population activity, a hysteresis like dynamics emerges. This novel phenomenon has important implications for determining the network response to time-varying inputs and for the network sensitivity at different operating points.

## Introduction

Neurons express a large diversity in terms of their biochemical, morphological and electrophysiological properties.^1–4^ However, it is not clear if and under which conditions such diversity plays a functional role. It has been shown that selective stimulation of neurons of a given type expressing specific bio-markers can modulate different aspects of brain functionn.^5^ For instance, selective stimulation of neurons changes the excitation/inhibition balance,^6^ network dynamics^7,8^ and computations performed by the network,^9^ thereby leading to an altered animal behaviour. Moreover, noise introduced by intrinsic properties of neurons/synapses can have several effects. It can render the dynamics more robust to perturbations^10^ and can improve the encoding and decoding of neuronal activity by reducing correlations.^11^ These experiments provide strong support to the ‘neuron doctrine’ and motivate the search for novel bio-markers and specific functions of different classes of neurons.^4,12^ However, experiments also suggest that stimulation of a certain neuron type may not cause any discernible change in the population activity and animal behaviour.^13^ Moreover, detailed models of single neurons^14^ and networks^15^ have shown that multiple combinations of neuron and synapse parameters can lead to similar activity states;^16^ suggesting that exact neuronal properties are not crucial to obtain a specific dynamical network state and, hence, a specific function.

These conflicting studies make it important to identify: (1) Changes in which of the neuron properties can affect the network dynamics? (2) In which dynamical states is the network activity susceptible to changes in a certain neuronal property? Here we focus on the effect of spike bursting on the network activity dynamics and vice versa. Spike bursting is a common electrophysiological descriptor of a neuron type^17,18^ and the fraction of bursting neurons depends on the brain region^19^ and even in a given brain region the firing rate of spike bursts may change depending on their inputs^20^ and on the behavioral task.^21^ Finally, the rate and fraction of burst spiking increases in epilepsy^22^ and Parkinson’s disease.^23^ From a dynamics perspective, when neurons operate in an ‘integration mode’, temporal integration of spike bursts can qualitatively change the response of post-synaptic neurons and, consequently, of the network. Such effects could be further amplified by short-term dynamics^24^ and long-term plasticity of the synapses.^25,26^ Therefore, the burst firing pattern, which is very different from the spike trains of the leaky-integrate-and-flre (LIF) neuron model is a suitable candidate to study the influence of single neuron firing patterns on the network activity. Surprisingly, despite this wealth of literature on the effects of neuronal and synaptic properties on network dynamics (see review by Wang^27^), it is not at all clear how firing patterns of various neuron types may affect the network dynamics and how network dynamics, in turn, may help shape neuronal firing patterns.

Here, we present an analytical framework to study the effect of spike bursting on the network dynamics. Using mathematical analysis and numerical simulations of large-scale network models of spiking neurons we investigate the effects of firing patterns - exemplified here by bursting activity of inhibitory neurons - on network synchrony and oscillations. Our analysis shows that there are two different mechanisms by which spike bursting can affect the network dynamics. We show that single-neuron burst firing is most effective in changing the network state when the latter is in a transition zone between asynchronous and synchronous firing regimes. That is, the effect of single-neuron bursting is contingent on the network activity state itself. Thus, our results suggest that the brain can exploit the heterogeneity of neuronal spike patterns if it operates in the transition zones between different activity regimes.

Finally, we show a novel property of hysteresis in the network activity, caused by the mutual interactions between single-neuron firing patterns and network dynamics. Hysteresis implies that the network output does not only depend on the current input but also on previous network states and that under certain conditions the network output will change slowly compared to the input. This will influence the network sensitivity at different operating points and, thereby, the network response to time-varying inputs.

## Results

Previous models have addressed the issue of neuronal and synaptic diversity by drawing values from various parameter distributions instead of assigning single values. The specific effect of neuronal heterogeneities in random networks becomes more apparent when instead of a distribution of neuron parameters, different types of neurons are used.^28^ Therefore, to study the effect of spike patterns of individual neurons, we characterised the activity of a randomly connected network of excitatory (*E*) and inhibitory (*I*) neurons by systematically increasing the fraction of one type of neuron in the inhibitory population. This manipulation was motivated by two experimental observations: (1) the fraction of bursting neurons depends on the brain region,^19^ (2) the probability of a neuron to elicit spike bursts depends on the inputs^20^ and neurons can dynamically switch their firing mode, depending on the context^21,29^ and, more permanently, in the case of specific brain diseases.^22,23^ That is, the fraction of bursting neurons is a dynamical variable which may change, depending on the behavioral context, inputs, brain region and brain condition. We considered a sparse Erdos-Renyi (ER) type network of *E* and *I* neurons connected with 10% probability. This choice of ER type random networks ensured that our results are not dependent on any specific connectivity of the bursting neurons. We used the Izhikevich neuron model for its computational efficiency and its ability to reproduce nearly all spike patterns observed *in vitro.*^30^ All excitatory neurons were realised as regular spiking neurons. The inhibitory neuron population consisted of *F%* burst spiking neurons (BS) and (100 – *F*)% fast spiking (FS) neurons.

### Effect of bursting on the stability of oscillatory activity

We first characterised the effect of bursting neurons on *γ*band oscillations in recurrent networks. These oscillations are considered to play a crucial role in brain function.^31–33^ We tuned the parameters - external input rate and synaptic weights - of a network of RS excitatory and FS inhibitory neurons (i.e. *F* = 0) to obtain stable *γ*band oscillations^34,35^ (Fig. 1B). In this regime, individual neurons do not produce an action potential in every oscillation cycle and, thus, have a mean discharge rate that is typically lower than the frequency of the fast gamma rhythm emerging at the network level.^36^ These oscillations are known to be robust to heterogeneities (when modeled by a unimodal distribution of neuron parameters) and noise in the external input.^36–38^ In the following, we study the stability of these oscillations in a network with two or three different types of neurons.

**Figure 1.**
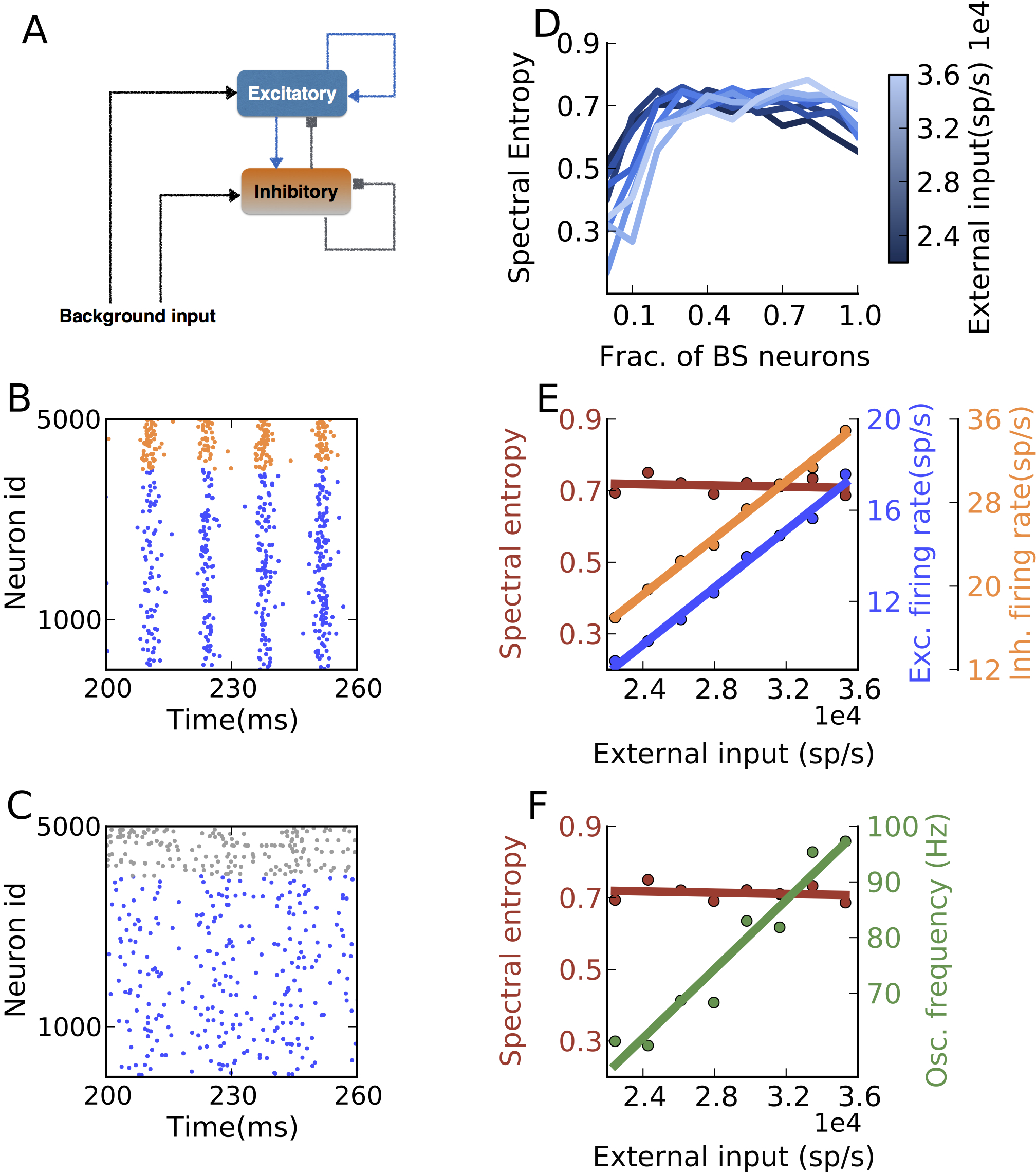
Effect of increasing the fraction of bursting neurons in the inhibitory population on the stability of *γ*band oscillations. (**A**) Schematic of the network. (**B**) Spiking activity in a network with only FS neurons constituting the inhibitory population. A clear oscillatory activity is seen in the excitatory neurons (blue dots) and inhibitory FS neurons (orange dots) (*g* = 7.1, *η* = 2.8 × 10^4^*sp/s, J_E_* = 0.1 *mV*). (**C**) Spiking activity in a network with only BS neurons (gray dots) constituting the inhibitory population. All other network parameters are the same as for the activity shown in B. Inhibitory BS neurons weaken network oscillations. (**D**) Stability of the oscillations (quantified by the Spectral Entropy) of excitatory neurons as a function of the fraction of BS neurons.(*g* = 7.1) (**E**) Spectral entropy, excitatory and inhibitory (FS+BS) population firing rate as a function of the external input (*η*) to a network with 40% BS and 60% FS inhibitory neurons. (F) Oscillation frequencies as a function of the external input. For a fixed fraction of BS neurons, spectral entropy remained unchanged while the oscillation frequency and the firing rate of the neurons increased.

When all inhibitory FS neurons were replaced by BS neurons, with all other parameters kept constant as in (Fig. 1B), the oscillations were severely weakened (Fig. 1C). For an intermediate fraction of BS neurons (*F* = 0.2), the oscillations were not completely diminished, but the stability of the oscillations was severely affected and short oscillatory epochs were interrupted by non-oscillatory activity. To quantify the stability of the oscillations, we estimated the spectral entropy (*H_s_*) of the population activity spectrum, which provides a measure of the dispersion of the spectral energy of a signal (see Methods). We found that the spectral entropy increased with the fraction of BS neurons and saturated to its maximum value (Fig. 1D). Irrespective of the strength of the external input (*η*), about 30% BS neurons were sufficient to quench the oscillations (Fig. 1D).

For a fixed proportion of BS and FS neurons, the excitatory input strength *η* shifted the operating point of the network by increasing the firing rate of the individual neurons (Fig. 1E). This also resulted in an increase in the dominant oscillation frequency (60 – 100 Hz), however, the spectral entropy remained unaffected (Fig. 1F). Thus, it is likely that the reduction in oscillation power is a consequence of the spike pattern of the BS and not of the different *f – I* curve of the BS neurons. Unfortunately, though, it is not trivial to separate the contribution of the spike patterns and the *f – I* curve to the network activity state. As we will show later, the effect of spike patterns and *f – I* curve can be separated by adapting the standard LIF neurons.

### Response of network activity to single neuron bursting

In the above, we showed the effect of BS neurons on the oscillatory dynamics of a random network only for a specific activity regime of the network. Sparsely connected random networks of excitatory and inhibitory neurons can exhibit distinct activity states depending on the external excitatory input (*η*) and the ratio of recurrent inhibition and excitation (*g*). While individual neurons can fire in a regular (R) or irregular (I) manner, the population activity can be synchronous (S) or asynchronous (A). Thus, the network activity could be either AI, SI, AR, or SR.^39,40^ In the mean-driven regime the neurons fire in a regular manner whereas in the fluctuation driven regime their spiking becomes irregular. Because neuronal activity *in vivo* is irregular, only SI and AI are biologically relevant for information processing. Therefore, we studied how the AI and SI activity regions in the parameters space of *η* and *g* are changed when FS are systematically replaced by BS neurons (Fig. 2A-C). The parameters *η* and *g* were varied to obtain low to mid-range firing rates ( ≤ 25spikes/sec) and irregular spiking in the RS neurons (CV_ISI_ ≥ 0.5).

**Figure 2.**
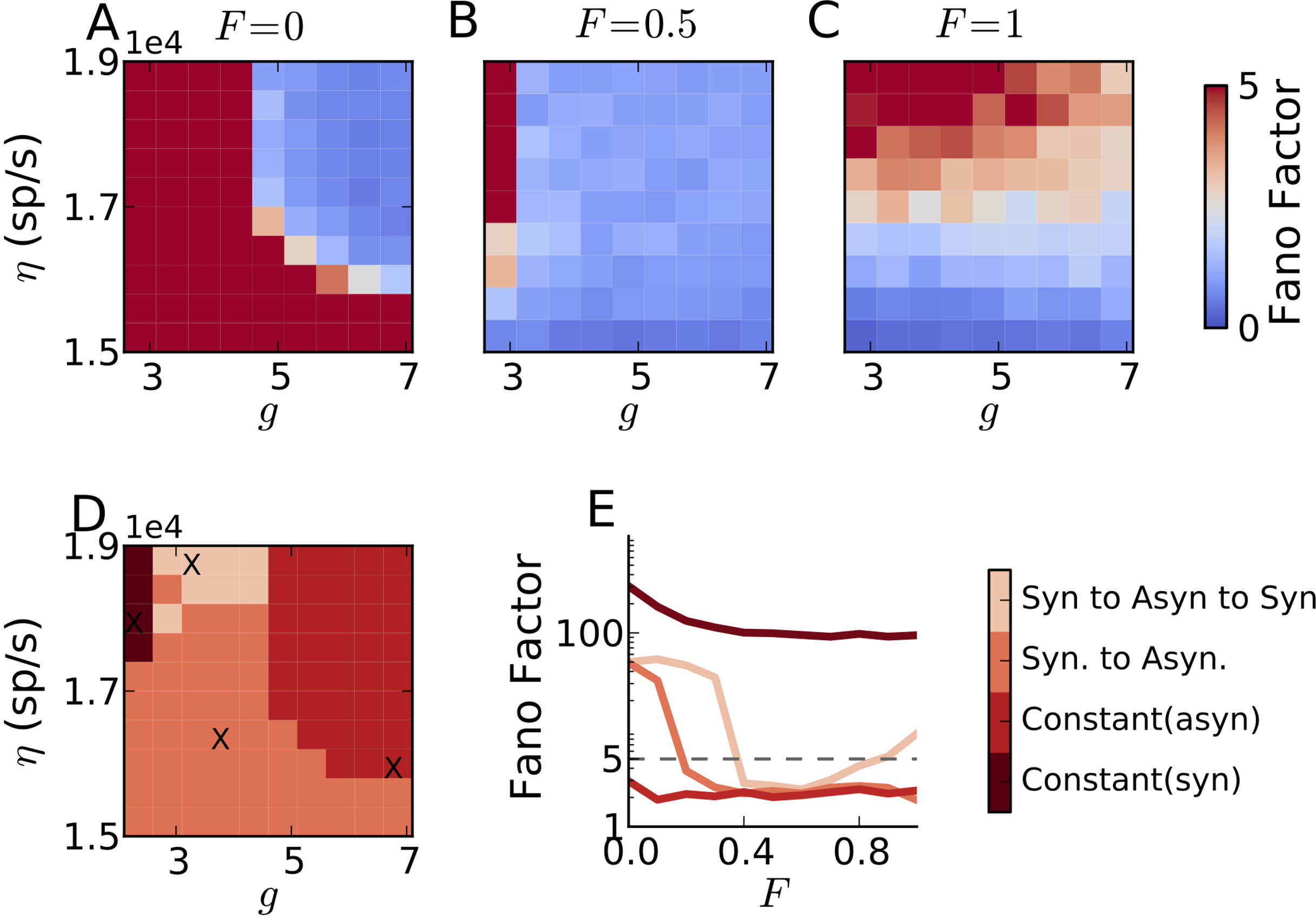
Effect of increasing the fraction of bursting neurons(*F*) in the inhibitory population on synchrony in the network activity. (**A**) Synchrony (measured as Fano Factor) in the excitatory neurons as a function of the ratio of recurrent inhibition and excitation (*g*) and external excitatory input (*η*), for 0% bursting neurons in the inhibitory population. (**B**) Same as in **A** when 50% inhibitory neurons are bursting type. (**C**) Same as in **A** when all inhibitory neurons are bursting type. (**D**) Summary of the changes induced by increasing fraction of bursting neurons on the different activity states of the network. (**E**) Four representative changes in the network synchrony as the fraction of bursting neurons is increased from 0 to 100% corresponding to the crosses in D.(*J_E_* = 0.1 *mV,d* = 1.5*ms*)

Replacement of FS neurons by BS neurons altered the various regions in the network parameter space differently. We identified four different ranges of parameters giving rise to four distinct modulations of activity regimes (see Fig. 2D,E): (1) A parameter range in which the network remained in the synchronous state, irrespective of the fraction of BS neurons. This invariance of the synchronous network activity to the neuron types was observed for small values of *g*. In a network where all neurons have identical *f − I* curve, this parameter regime would correspond to a mean-driven regime. This classification is, however, not directly applicable here, because FS and RS neurons have different slopes of their *f − I* curves. (2) A parameter range in which the network remains in an asynchronous state, irrespective of the fraction of BS neurons. In this regime, *g* is large enough to drive the network into the fluctuation-drive state, resulting in irregular and asynchronous (non-oscillatory – *H_S_ ≥* 0.6) firing. (3) The network activity makes a transition from the synchronous to the asynchronous state, that is, BS neurons tend to weaken or even quench the weak synchrony. (4) In a relatively small parameter regime, we also observed that for a small fraction of BS neurons, the network activity changed from the synchronous to the asynchronous state (similar to (3)), but for a larger fraction of BS neurons, the activity returned to the synchronous state again. That is, for 100% BS or FS neurons, the network remained in a synchronous (also oscillatory, *H_S_ ≤* 0.6) state, whereas for intermediate fractions the network synchrony was destroyed(*H_S_* ≥ 0.6).

The Izhikevich neuron in its bursting mode, differs from its fast-spiking mode in two respects: it produces more than one spike every time the membrane potential crosses the spiking threshold (see Fig. 3A) and the *f — I* curve of the bursting neurons has a larger slope than that of the FS neurons (see Fig. 3B). In the existing neuron models (Izhikevich neuron model, generalised integrate-and-fire neuron), it is not possible to change the *f − I* curve of the neuron without affecting its firing pattern.

**Figure 3.**
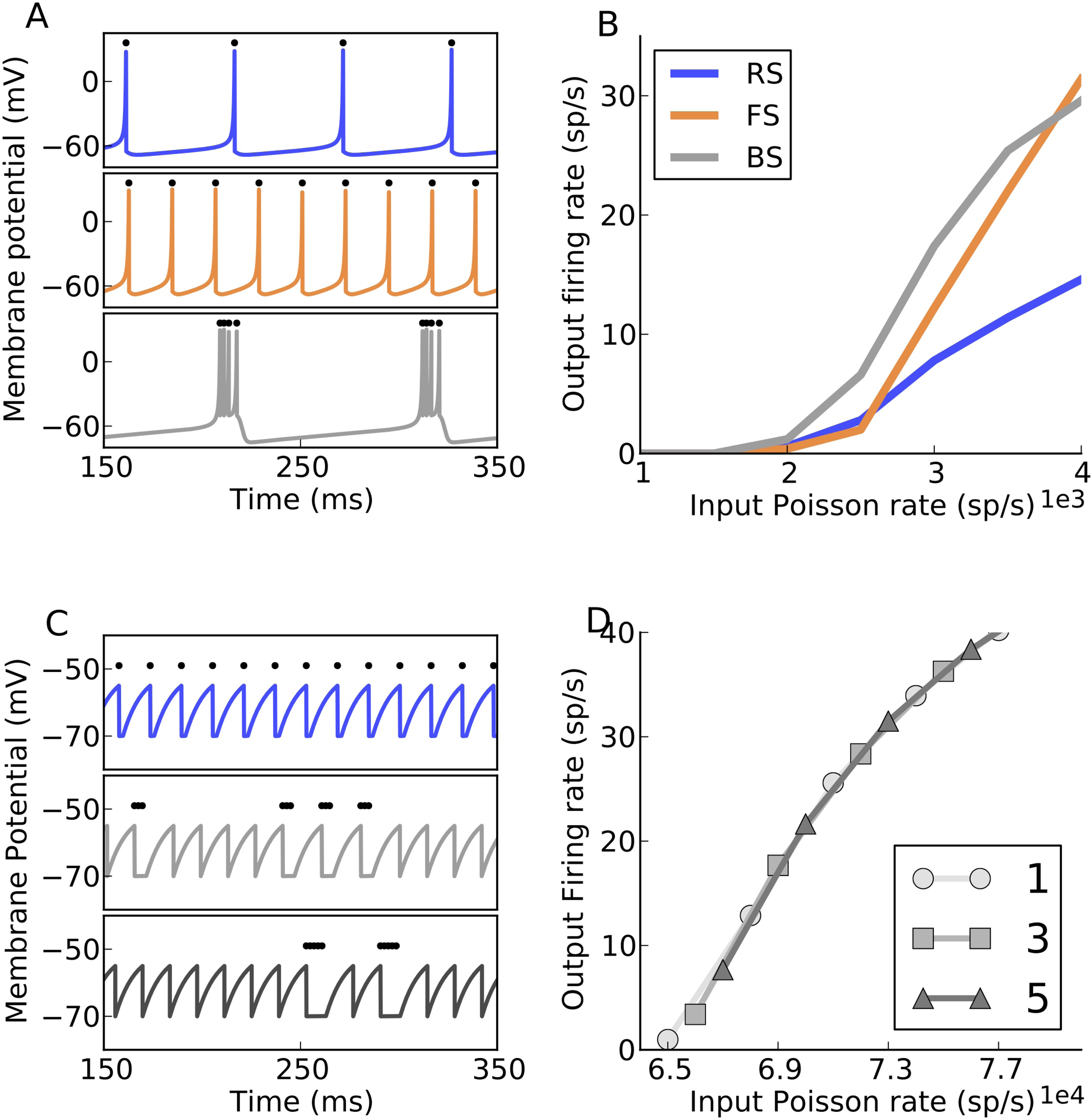
The state-dependent stochastic bursting neuron. **(A)** Izhikevich neuron producing regular spiking (RS), fast spiking (FS) and Bursting (BS) firing patterns for different values of the neuron parameters a,b,c and d. **(B)** The firing rate response of the neuron types for different poisson input rates. **(C)** Firing patterns of the State-dependent Stochastic Bursting Neuron (SSBN) model with varying number of spikes per burst for the same value of constant external DC input (top to bottom). **(D)** The firing rate response curve of the SSBN for different number of spikes per burst, for external Poisson input of different rates.

### The state-dependent stochastic bursting neuron model

To understand the role of spike patterns in shaping the network dynamics it is important to isolate their effects from the different *f − I* curves. Therefore, we modified the standard LIF neuron model to produce bursting of *B* spikes in a stochastic manner with a probability 1/*B* every time its membrane potential reaches the spiking threshold (see Methods). We refer to this new model as the State-dependent Stochastic Bursting Neuron (SSBN) model when the parameter *B* depended on the input level. In a special case, *B* could be a fixed number. The SSBN model not only ensures that the *f − I* curves of the bursting and fast-spiking neurons remain identical (Fig. 3D), but it also allows us to change the size and the duration of the burst without cumbersome parameter tuning (Fig. 3C). Moreover, unlike the Izhikevich neuron model and the generalised LIF model, which are often used to model bursting dynamics of neurons, the bursting characteristics of the SSBN remain unchanged, irrespective of the input statistics. The response characteristics of the SSBN are similar to that of the LIF, except that an increase in the number of spikes per burst B decreases the high-frequency firing limit of the neuron (Supplementary Fig. S1).

### Effects of different firing patterns of inhibitory neurons on the stability of network oscillations

In contrast to FS neurons, BS neurons spike in bursts, but for the same input the total number of spikes generated by a BS neuron is identical to that of an FS neuron. This implies that in the SSB neuron, spikes are clumped together, creating ‘empty’ temporal windows (with a duration depending on burst size) in which no spikes occur (Fig. 3C) and very short windows in which the number of spikes produced will be significantly higher than that of FS neurons. Therefore, while an FS neuron exerts a relatively uniform inhibition onto its post-synaptic neurons, BS neurons exert inhibition in clumps. Because of the temporal clustering of spikes in BS neurons, two distinct mechanisms emerge that define the stability of the oscillatory and asynchronous states, respectively.

### Stability of the oscillatory state: Additional spikes part of the burst disrupt oscillations

Fast (or γ) oscillations could be described as ‘interneuron generated’ (ING) or pyramidal-interneuron generated (PING).^37^ In the ING oscillations, recurrent inhibition of the inhibitory interneurons creates a small time window for pyramidal neurons to spike. In the PING mechanism, an increased activity of pyramidal neurons causes an increase in the activity of inhibitory interneurons, which subsequently inhibit the pyramidal neurons. In both mechanisms, inhibition sets the time window for the activation (ING) or inactivation (PING) of the pyramidal neurons.^41^ The temporal clustering of spikes in BS neurons causes a temporal jitter in the duration of the recurrent inhibition and, therefore, weakens the oscillations (mechanism-I).

This is best illustrated in the case of ING oscillations. Here, the initiation of a burst at the edge of the preceding oscillation cycle distorts the subsequent window of opportunity for the next inhibitory cycle and, consequently, the oscillation is quenched in the inhibitory population. This renders the excitatory population non-oscillatory as well.

To demonstrate this mechanism, we simulated a simple E-I network with an inhibitory population composed of FS neurons only. The values of *g* and *η* were adjusted to render the network in the ING oscillation regime. Based on thresholding the Z-scored PSTH of the population activity, the oscillatory cycles were marked (gray stripes in Fig. 4A). Next, we simulated the network once more with identical parameters, except that at the fifth oscillatory cycle (Fig. 4A), 40% of FS neurons were replaced by BS neurons. By comparing these two simulations, we determined the number of ‘additional’ inhibitory spikes (*num_add_*) that fell outside the oscillatory window.

**Figure 4.**
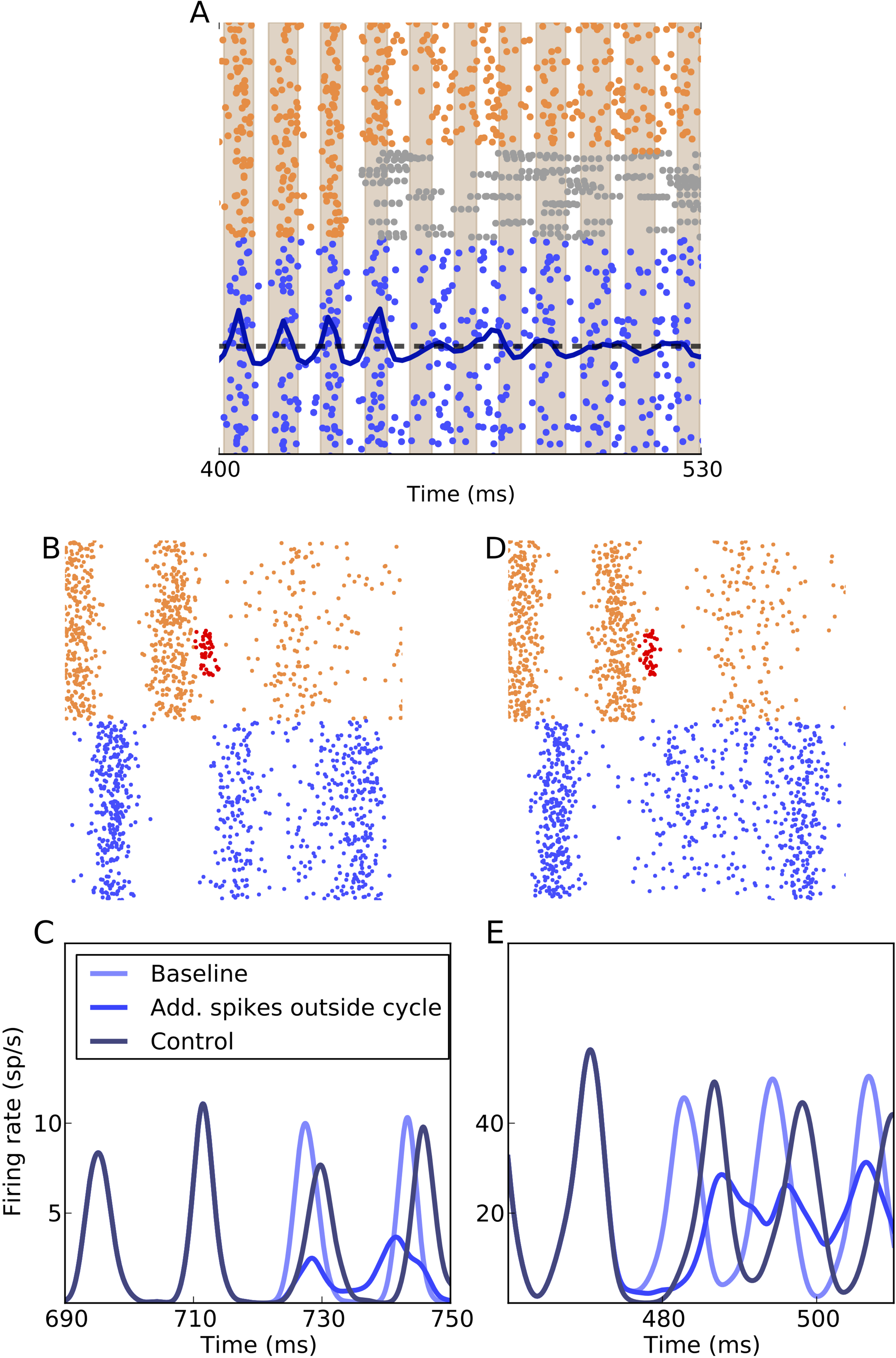
Mechanism-I by which spike bursting destroys network oscillations.(**A**) The network which is initially in an oscillatory state switches to a non-oscillatory state with the replacement of FS neurons(orange dots) with the BS neurons (grey dots) in the inhibitory population. The blue dots show the excitatory spikes and the dark blue line is the z-scored PSTH of the excitatory activity. The light brown stripes correspond to the crest of the oscillatory cycles of the excitatory population when the network consisted of only FS inhibitory neurons. The number of additional spikes that fall within the stripes is calculated(*num_add_*) added (*g* = 12*,d* = 2*ms,η* = 11500*sp/s,F* = 0.4)(**B**) A schematic to depict how the addition of additional inhibitory spikes(red dots) when the inhibitory oscillatory cycle wanes makes the oscillatory activity unstable in an ING oscillation. The excitatory population (blue dots) oscillates in the window of opportunity provided by the inhibitory population (orange dots). The red dots indicate the additional inhibitory spikes that are added. (**D**) Same as in (C), except that the oscillations are PING driven. (**D**) PSTHs of the excitatory population shows the changes after the addition of the *num_add_* additional spikes in the inhibitory population. When the additional spikes are added when the inhibitory oscillatory cycle tapers off there is maximum disturbance of succeeding oscillatory cycles (blue line). When the same number of spikes are added at the peak of the preceding oscillatory cycle, there is minimal effect on the subsequent oscillatory cycle (dark blue line). The pale blue line shows the baseline activity when no spikes are added (g = 12*,d* = 2*ms, η* = 11500*sp/s*). (**E**) PSTHs of the excitatory population affected by additional spikes in a PING driven oscillation(*g* = 7*,d* = 1.5 *ms, η* = 20000 *sp/s*).

To mimic the effect caused by the additional spikes generated by the BS neurons, we added *num_add_* additional inhibitory spikes at the exact moment when a particular inhibitory oscillatory cycle tapered off (Fig. 4B). This time was determined by running an identical simulation with the same random number generator seeds (baseline) (Fig. 4C pale blue trace). Addition of the previously determined number of extra inhibitory spikes (as would happen in a BS neuron) indeed disturbed the next oscillatory cycle significantly (Fig. 4C blue trace).

To test whether it is the timing of the bursts that weakens the oscillations and not the number of spikes contained in them, we added the same number of additional inhibitory spikes during the peak of the preceding inhibitory oscillatory cycle (control). In this case, the oscillation amplitude and frequency were not significantly changed (Fig. 4C dark blue trace), thereby showing that only the timing of the bursts (or the corresponding additional spikes) destroyed oscillations. A similar distortion of oscillations is observed when adding additional spikes in the inhibitory population in a network in which oscillations are driven by the PING mechanism (Fig. 4E) (scheme in Fig. 4D). The breakdown of oscillations by temporal jitter of inhibition is effective when oscillations are weak. In strongly oscillatory states, the effective synaptic couplings are strong and, hence, jittering of inhibition is not sufficient for quenching oscillations (see also Supplementary Fig. S2B).

### Stability of the asynchronous state: Bursting makes the network susceptible to oscillations

When spikes arrive in a burst, the post-synaptic neuron receives a much bigger compound PSP due to the temporal summation of individual spikes. Because we preserved the *f − I* curve of the neuron while making it bursting, effectively each spike was replaced by *B* spikes while reducing the input rate by a factor *B*. This is equivalent to a network of non-bursting neurons connected with a synaptic kernel that reflects the temporal summation of spikes in the burst. This analogy allows us to use the established mean-field theory to investigate the stability of the AI state of the network activity.^36,38^ Only when the compound PSP renders the AI state to become unstable, we would expect bursting neurons to transform the AI state into the SI state, otherwise a change in the neuron spiking behavior will not affect the network activity.

For simplicity in our network we kept the recurrent synaptic coupling strengths as *J_EE_* = *J_IE_* = *J_E_* and *J_II_* = *J_EI_* = *J_I_*, and *J_I_* = *g* · *J_E_* (where the subscript xy indicates a connection from the *y* population to the *x* population). To check for the stability of the AI state, we introduced a small perturbation in the steady-state firing rate 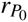 of population *P* (excitatory or inhibitory),

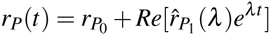

where *λ = x + jω* with *ω* being the modulation frequency. The perturbation in the steady-state firing rate leads to a perturbation in the recurrent synaptic input

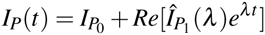

where 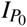 is the baseline steady state synaptic input, 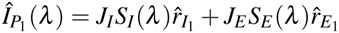, an d *S_I_* and *S_E_* are the synaptic response functions.^38^

Subsequently, the perturbation in the synaptic input would change the network firing rate by 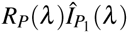 (where *R_p_*(*λ*) is the neuron response function^38^). In a recurrent network, if the rate perturbation, 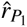 is equal to the synaptic input perturbation, the perturbation does not die out, indicating an instability of the asynchronous state. That is, for an unstable asynchronous state:

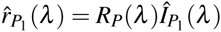

We used the above equation to derive the conditions for the instability of the AI state by analyzing the following equation:^38^

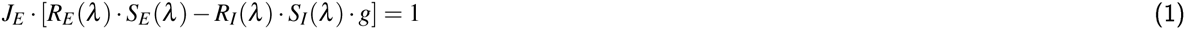

where 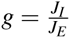. If the synaptic coupling strength *J_E_* crosses a critical *J_cr_*, the asynchronous activity destabilizes and the network activity enters an oscillatory regime. Because of the temporal summation of burst spikes, when BS neurons replace FS neurons in the inhibitory population, the inhibitory synaptic response function *S_I_* is altered. Specifically, an increase in the number of spikes per burst leads to an increase in the effective synaptic rise time (see Methods). This in turn, leads to a reduction of the critical coupling value *J_cr_*, rendering the AI state unstable (see Fig. 5A – black dotted line). Thus, if *J_E_* < J_cr_ for *B* = 1 and *J_E_* > *J_cr_* for *B* = 4, a change of neuron type from FS (*B* = 1) to BS (*B* = 4) will destabilize the AI state and lead the network activity into an oscillatory state. However, if *J_E_* remains below *J_cr_* for *B* = 1 and *B*=4 the network remains in the asynchronous state, despite the replacement of FS by BS neurons. If the network with FS neurons is already in a synchronous state (*J_E_* > *J_cr_*), a replacement of all of the FS neurons with BS neurons will not affect the state. However, if the oscillations are weak, replacement of a certain fraction of FS neurons with BS neurons can destroy oscillations through mechanism-I by temporal jitter of inhibition. Thus, in the asynchronous activity state BS neurons affect the network dynamics by reducing the value of the critical coupling (*J_cr_*), leading to a shift from asynchronous to synchronous network activity (mechanism-II). As equation-1 indicates, whether or not BS neurons will change the asynchronous activity state to the oscillatory state by mechanism-II depends on the network connectivity parameters and the firing rate of the network *r*_0_.

**Figure 5.**
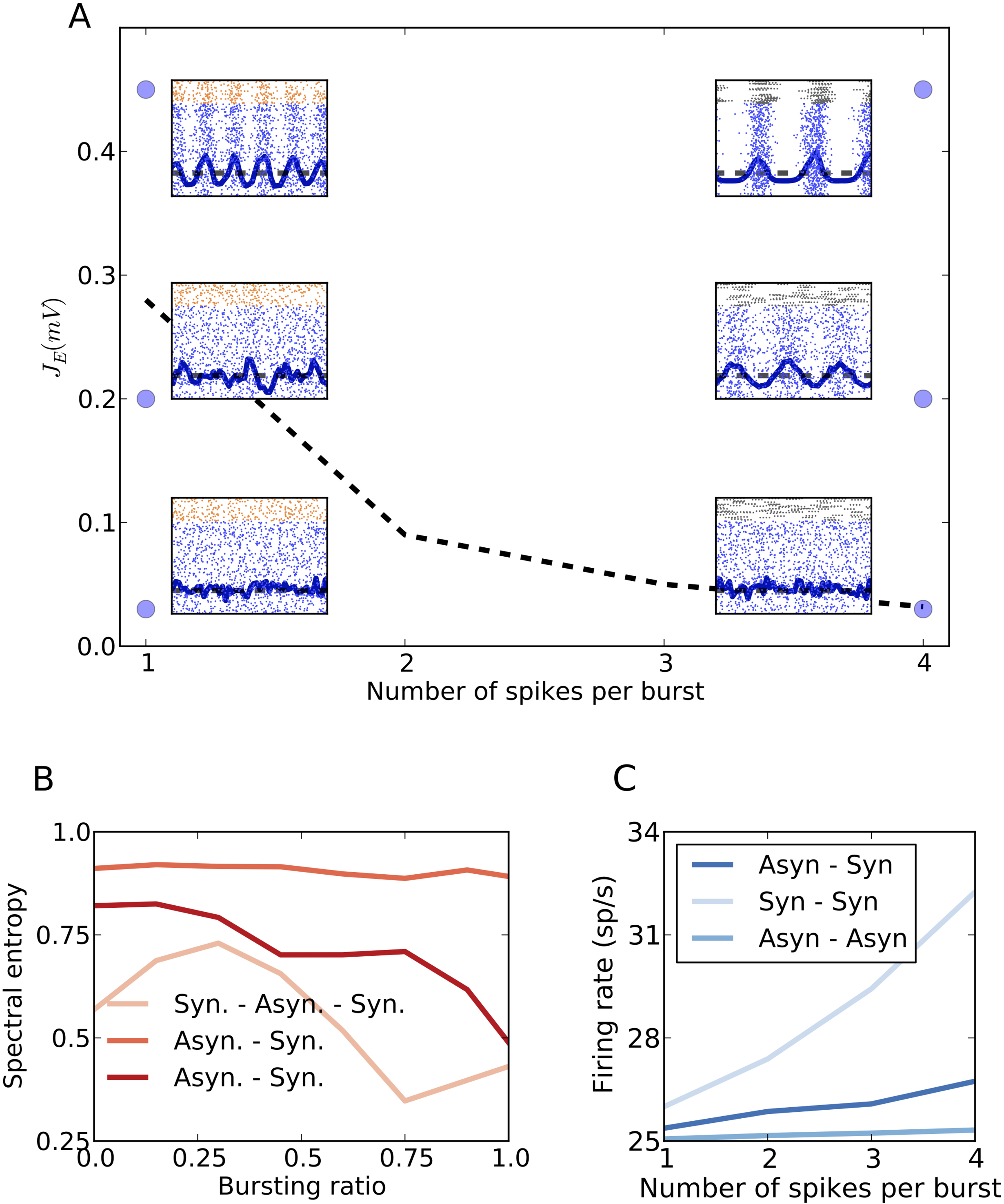
Mechanism-II by which spike bursting enhances oscillations. **(A)** In the phase space of excitatory synaptic strength (*J_E_*) and the number of spikes per burst(*F*), the bifurcation line (dotted black line) between the oscillatory and non-oscillatory states is the *J_cr_* value calculated analytically (for input *mean* = 14*mV* and *σ* = 6*mV*. *d* = 5*ms, t_syn_* = 1 *ms* and *V_th_* = 20*mV).*. When the FS neurons in the inhibitory population are replaced by BS neurons the number of spikes per burst of the neurons in the inhibitory population is altered and the *J_cr_* value drops. A network in an initially asynchronous state can continue to remain asynchronous with the addition of BS neurons if the *J_E_* values are less than *J_cr_* for *F* = 4 (bottom panels). The network can transition from asynchronous to synchronous states with the change in F, if the *J_E_* is more than *J_cr_* for *F* = 4 (middle panels). Also, a network in an oscillatory state for *F* = 1 remains oscillatory for *F* = 4 (top panels). (B) Instead of replacing the entire FS population with BS neurons, different proportions of the inhibitory population were changed for the networks in panel **A** with *F* = 1. It is observed that the addition of 25% BS neurons in a network in a synchronous state, destroys oscillations due to Mechanism -I. **(C)** The change in the firing rate of the excitatory population for transitions in **A** while number of spikes per burst are changed.

### Effect of spike bursting on the network activity dynamics

The understanding of how BS neurons or spike bursts affect the network dynamics allowed us to re-examine the change in the dynamics of a recurrent network when FS neurons are systematically replaced by BS neurons. We simulated a random recurrent network with SSB neurons and studied the robustness of the synchronous and asynchronous states when single spiking SSB neurons (equivalent to FS neurons) were systematically replaced by SSB neurons with spike bursts of size four (equivalent to BS neurons).

When the network was tuned to be in an oscillatory regime (*J_E_* > *J_cr_*), an increase of the number of bursting neurons first lead to a non-oscillatory network activity (*H_s_ ≈* 0.75, *F* = 25%). This weakening of the oscillations is a result of mechanism-I. However, as the fraction of BS neurons was further increased (F ≥ 50%), mechanism-II became more effective and counteracted mechanism-I, resulting in oscillatory network activity again (*H_s_ ≈* 0.5) (see Fig. 5B). This non-monotonic change in *H_s_* resembles the non-monotonic change in networks with the Izhikevich model neuron (see Fig. 1A-C). Based on our observations made in networks with SSB neurons, we think that even in a network with Izhikevich model neurons, the non-monotonic state changes were largely governed by the change in neuron spike patterns. Note that a network can remain in the synchronous state for all values of *F*, provided that the inputs to the excitatory and inhibitory populations are appropriately controlled (see Supplementary Fig. S2B).

When the network was tuned to be in an asynchronous non-oscillatory state with weak correlations (*H_S_ ≈* 0.7, *J_E_ < J_cr_*), replacing FS neurons by BS neurons rendered the network in an oscillatory state. The spectral entropy monotonically decreased with the fraction of BS neurons (see Fig. 5B). Hence, the transformation of non-oscillatory activity to the oscillatory state was governed purely by mechanism-II.

In a network with highly aperiodic activity and very weak correlations (*H_S_ ≥* 0.8, *J_E_ ≪ J_cr_*), i.e. when the activity is deep in the AI regime, the network state was robust to changes in the spike pattern properties of individual neurons (see Fig. 5A).

These results clearly show that neuron spike patterns can indeed change the network state, from a weakly non-oscillatory asynchronous state to synchronous oscillations (by mechanism-II) and vice versa (by mechanism-I). At the same time, a non-oscillatory state with very weak correlations is invariant to changes in the neuron spike pattern properties. We conclude that network activity is susceptible to neuron spiking patterns only in the transition zone between different regimes (here between asynchronous–non–oscillatory and synchronous–oscillatory) and the effect of neuron spike pattern properties on the network activity dynamics is contingent on the network activity state itself.

### Bursting activity increases the population firing rate

The bursting firing pattern of the inhibitory neurons aids in the transition of the network activity from the asynchronous to the synchronous state (Fig. 5A). This change in the stability of the network activity also influences the population firing rate (Fig. 6). The increasing ‘burstiness’ of the constituent bursting neurons steers the network activity into an oscillatory state. This switch is accompanied by an increase in the population firing rate.

**Figure 6.**
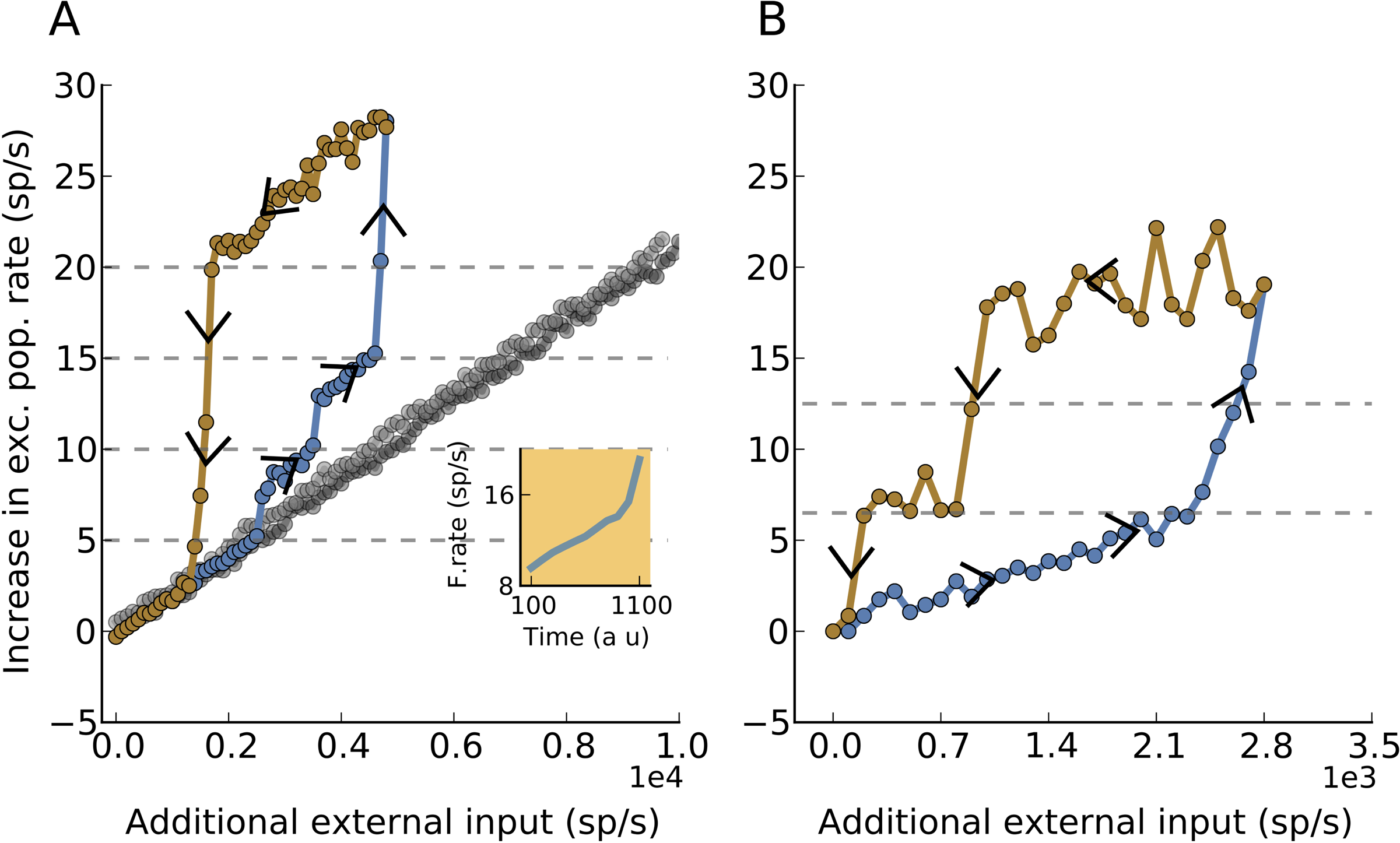
Bursting introduces multi-stability and hysteresis in the network dynamics. (**A**)The increase in firing rate due to increase in external input and change in the burstiness of the neuron (dashed grey lines) is shown. The simulation protocol to generate this neuronal network hysteresis is described in Methods. It is seen that the onward (blue line) and return (brown line) curves do not trace the same path indicating the state dependence of the effect of the single neuron firing pattern on the network. The grey dots show a similar hysteresis loop for a network in which the burstiness of only 20% of the inhibitory neurons is changed. The inset plot shows the simultaneous change of the firing rate of the network and the burstiness of the modified-SSBN after given an initial perturbation of additional external input of 200 spikes/sec. The burstiness of the inhibitory neurons (as defined by the state variable (see Methods)) increases with the excitatory population firing rate.The increase in bursting in turn increases the population firing rate. This self-propelling mechanism continues till the single neurons produce the maximum number of spikes per burst (*B* = 5).(**B**) This panel is similar to **A**, but the firing rate estimate of the excitatory population is made over a time window of 200*ms*. The number of spikes per burst increase by 2 for every crossing of the firing rate threshold.

Additionally, the difference in the temporal structure of bursting could also change the statistics of the total synaptic inputs and the output firing rate of a postsynaptic neuron. To test this, we fixed the number of bursts of an SSB neuron and connected it to a LIF neuron that also received excitatory Poisson input. We measured the output firing rate and variance of the free membrane potential *v_fr_* of the post-synaptic LIF neuron as a function of the number of spikes in a burst (Supplementary Fig. S3). The mean *v_fr_* remained constant as the number of spikes in a burst was increased, because irrespective of burst size the total numbers of excitatory and inhibitory spikes were preserved. However, the temporal clustering of BS spiking increased the variance of *v_fr_*, resulting also in an increase of the output firing. At the network level, this could also contribute to an increase in the population firing rate, thereby reducing *J_cr_* and, hence, contributing to the switch of activity from the asynchronous irregular to the oscillatory state by facilitating mechanism-II.

### State dependent bursting of inhibitory neurons induces hysteresis in the network dynamics

In the above, we made the assumption that the number of spikes in a burst of the SSB neuron is fixed. In real neurons, where spike bursting is governed by the voltage-dependent ion channels and interactions between soma and distant tufts (e.g. in pyramidal neurons^20^), the number of spikes in a burst would depend on the network activity level. Consistent with this, recent experiments indeed show that bursting can change, depending on the behavioural task and the network activity state^21^ in both excitatory and inhibitory cells. In simulations with networks of Izhikevich neurons, we found the ‘burstiness’ of BS neurons also to be dependent on the network activity state (Supplementary Fig. S4).

To implement such state-dependence of burst size, we quantized the firing rate of the excitatory neurons into five disjunct ranges (*l_B_* = (B-1)**×**5 − B×5 spikes/sec., with *B* ϵ {1,2,3,4,5}). The SSB neuron generated *B* spikes per burst, depending on the level of the firing rate of the excitatory neurons.

With this model of state-depending bursting in inhibitory neurons, we further explored the relationship between the network level and neuron level properties. Usually, stationary Poisson inputs are used to determine the steady state of the network activity. However, such steady state will not reveal any effects introduced by state-dependent bursting of inhibitory neurons. Here we introduced dynamical changes in the network activity by slowly varying the external input (100 spikes/sec per observation window 3sec or 200ms;see Methods).

Random recurrent network without any state-dependent changes in neuron properties rapidly follow changes in the external input^42^ (Fig.6A, black dots). By contrast, networks with SSBNs, exhibited hysteresis, that is, when the input was changed slowly, the response of the network depended not only on the current input value but also on its history (Fig.6A,B orange dots).

To understand the hysteresis observed here, it is important to recall that the change in the population firing rate in the system was determined by two factors: (1) a change in the external input, and (2) a change in the number of spikes per burst (*B*) of the SSB neurons. An increase in the external input rate led to an increase in the network population firing rate, until SSB neurons started to burst. Therefore, any further change in the network firing rate was governed by both the further rising input rate and the increasing effect of neuron bursting. Moreover, every time *B* was increased (see Methods), the network activity rapidly jumped (Fig. 6 A). At the peak network output firing rate, when the SSB neuron elicited 5 spikes per burst, the increase in the network firing rate was dominated by the increase in *B*. In this network state, a reduction of the external input had only a very weak effect in decreasing the population firing rate, until the network firing rate had dropped enough to reduce the burst size. Once the activity dropped below this range, it rapidly returned to the baseline state. In the case of a network with a small fraction of BS neurons (20%), the increase in network firing rate due to the change in *B* was very small (Fig. 6A black dots), resulting in very little difference between the network responses during the increasing and decreasing cycles of the external input.

Balanced random networks, which are often used to model cortical network activity, do not exhibit such hysteresis properties in biologically relevant activity regimes such as the asynchronous-irregular or synchronous-irregular states.^39,42^ However, under some special conditions, such as clustered connectivity^43^ and plastic synapses,^44,45^ spiking neuronal networks can exhibit bistability that may lead to hysteresis as well. Hysteresis in network activity implies slow dynamics. On the one hand, bursting increases the sensitivity of the network to slowly varying changes, but on the other hand, hysteresis could result in a persistent activity – that is, a change in network response activity, lasting long after the stimulus originally inducing it has passed.

## Discussion

A specific neuron type has a functional significance only if it has a discernible effect on the network activity state. At the level of spiking activity, the effect of neuronal parameters can be described in terms of changes in the firing pattern (e.g. bursting and non-bursting) and *f − I* curve (Fig. 7A). Here, we investigated when and how neuronal spike bursting, one of the most common descriptors of neuronal types, can introduce a qualitative change in network activity. Our theoretical analysis and numerical simulations of neuronal networks show that the impact of spike bursting is contingent on the network activity state (schematically shown in Fig. 7B). The change in the network activity state caused by the temporal clustering of spikes in BS neurons can be understood in terms of two mechanisms (Fig. 7A,B). When the network operates in a moderately oscillatory regime (spectral entropy ≈ 0.5), spike bursts distort the temporal relation between the excitation and inhibition necessary for these oscillations^37,41^ and, therefore, weaken the oscillations (mechanism-I). In this regime, BS neurons increase the noise, thereby weakening oscillations (Fig. 4, Fig. 7B). On the other hand, spike bursting reduces the effective coupling strength *J_cr_* (see eq. 1), causing the asynchronous activity state to destabilize (mechanism-II). That is, bursting reduces the region in the network parameter space for which asynchronous activity is stable (Fig. 5, Fig. 7B). These two mechanisms are most in effect when the network activity is in a region in the activity state space, close to the border between asynchronous and oscillatory states. By contrast, the highly asynchronous and fully synchronous states remain unaffected by the change in the neuron spiking behavior caused by ‘replacing’ FS neurons by BS neurons.

**Figure 7.**
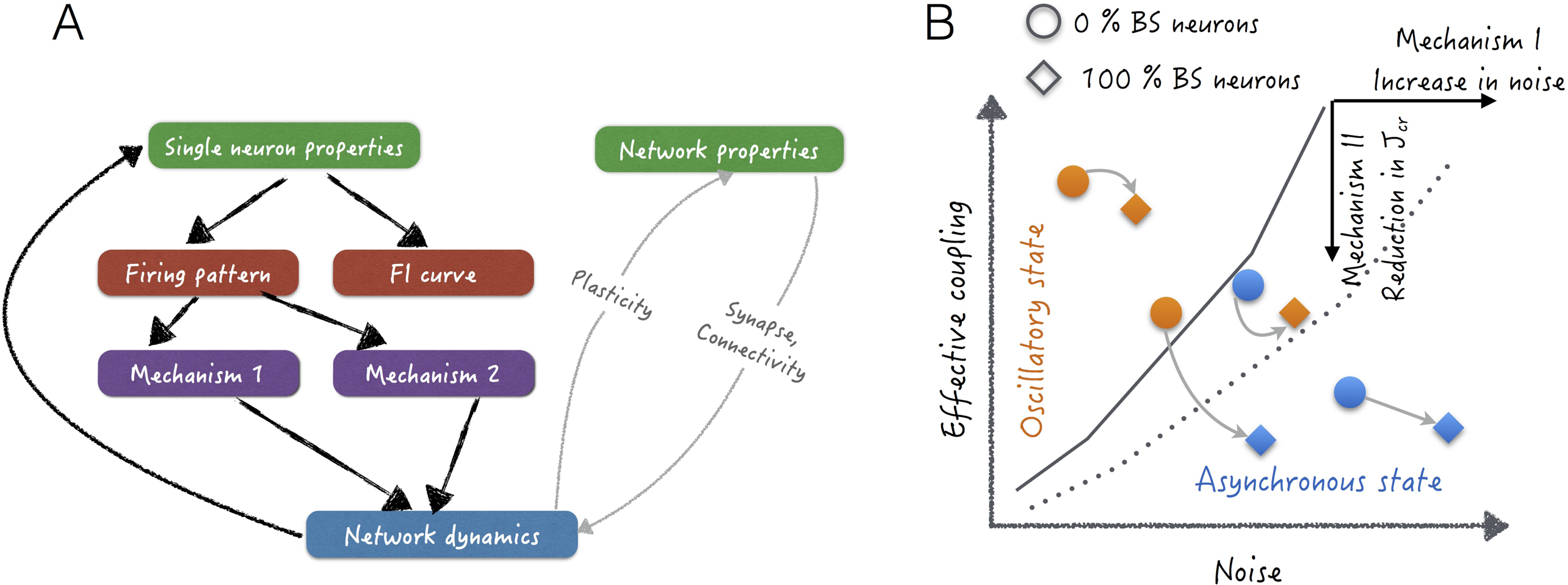
(**A**) The flowchart shows the different aspects that we glean from the results to establish the relationship between the single neuron properties and network dynamics (black lines). The description of the network effects of bursting through the two mechanisms was achieved by separating the effect of ƒ − *I* curves from that of the firing patterns by using SSB neurons. The single neuron firing pattern made dependent on the network dynamics resulted in hysteresis. The gray lines show the unexplored facets of the relationship between the two in the manuscript.(**B**) This schematic summarizes how the two mechanisms control the oscillatory activity in the network. The addition of BS neurons in an oscillating system gives rise to a recurrent noise and destroys the fine temporal balance between E and I populations that give rise to oscillations and quench them. Mechanism-II shifts the bifurcation line in the phase space by reducing the *J_cr_* with the addition of BS neurons.

### Functional consequences of a bursting dependent network state change

We showed that weak oscillatory activity is especially susceptible to spike bursting and that even a low fraction of BS neurons (*≈* 30%) in the inhibitory population is sufficient to quench oscillations (mechanism-I). Such a transient increase in the activity of BS neurons could form a powerful mechanism to reset network oscillations. Network oscillations in γ band (30-80Hz) are considered to form the basis of selective communication between weakly connected brain regions.^32,41^ Bursting-induced phase resetting could be a powerful mechanism to stop or start a communication between two such brain regions. Recent experiments show that bursting does indeed increase in a task-dependent manner and that it synchronizes activity between different brain areas.^21^ Our results provide two potential mechanisms that can act to induce phase-resetting and/or phase-synchronization and, therefore, provide a first theoretical account for these experimental findings.

In our study, we did not incorporate any specific connectivity of the bursting neurons and, therefore, may have underestimated the effect of spike bursting on the network dynamics. Recent experimental data suggest that neurons exhibiting different firing patterns may receive inputs from different sources.^46^ Given that neuronal connectivity is a key determinant of the effect a given a neuron has on the overall network dynamics,^47–50^ the effects of spike bursting on network activity would be further accentuated when bursting neurons make more specific connections, which might possibly form in networks with activity dependent synaptic plasticity (Fig. 7A).

### Network hysteresis

Spike bursting could be an intrinsic property of neurons^51^ or emerge as a consequence of network activity.^20,21^ In our simulations, when we made the burst size dependent on the average firing rate in the network, we observed a hysteresis-like behaviour for time-varying inputs (Fig. 6). Classical balanced random networks closely track the dynamics of the external input and do not show such behaviour - in fact, a hallmark of their behaviour is to track an arbitrarily fast external input.^42^ Interestingly, the speeding up or slowing down of network dynamics due to the presence of bursting neurons has also been observed in other complex networks with bursting communication patterns for specific network configurations.^52^

To the best of our knowledge this is the first demonstration of hysteresis in Erdos-Renyi random recurrent network models of cortical networks, with weak static synapses and sparse connectivity.^27,39^ Typically, in network models, low-level neuron and synapse properties affect network dynamics and not the reverse, as we have shown here. Notable exceptions are networks with plastic synapses^53^ and conductance-based synapses.^40^ Hence, we suggest that searching for hysteresis-like behaviour in experiments could be a promising approach to identify mutually causal influences between low-level neuron properties and high-level network dynamics.

When the size of the spike burst and the network activity are mutually dependent, the network gain depends both on the network activity state and the history of the input. This is quite unlike the conventional balanced random networks, where the input history plays no role in determining the network gain. More work is needed to fully understand how such input-history-dependent changes in the network gain will affect the processing of time-varying input signals.

Finally, we speculate that disease-related aberrant neuronal activity could be a consequence of an increased fraction of bursting neurons, e.g. in Parkinson’s disease^23^ and epilepsy.^22^ In these cases, possible treatments could aim at identifying and counteracting the precise mechanisms of bursting activity, either pharmacologically or through electrical stimulation.

### Conclusions

In summary, bursting neurons may play a crucial role in coordinating communication between different brain areas, by affecting the oscillation phase of network oscillations, they may induce hysteresis and, thereby, persistent activity in the networks, and they could even alter the global activity state of the network. From this, it is evident that single neuron properties have a significant impact on network dynamics, but this is possibly only the case in certain network activity regimes. Therefore, the effects of low level neuron and synaptic properties can be understood only in the context of higher level network activity attributes. This complex interplay between low and high level features introduces emergent phenomena that enrich the dynamical repertoire of the brain.

## Materials and Methods

### Neurons

#### Neuron model

Here we used the phenomenological model introduced by Izhekevich.^30^ The sub-threshold dynamics of this neuron model is defined by

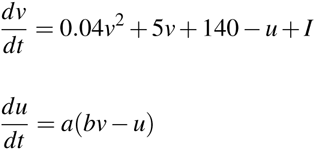

and the spiking is described by if *v ≥* 30 mV, then *v ← c* and u *← u + d*

The variable *v* denotes the membrane potential and *u* denotes the activation of K^+^ ionic current and inactivation of Na^2+^ ionic current. The parameter *a* determines the time scale of the recovery variable and *b* defines the sensitivity of *u* to the subthreshold fluctuations of *v*. *c* and *d* determine the reset values of *v* and *u* after spiking respectively. The parameters used for the three types of neurons are given in **Table 1.**

**Table 1.** Izhikevich neuron parameters

| Neuron type | a | b | c | d |
| --- | --- | --- | --- | --- |
| Regular spiking(RS) | 0.2 | 0.2 | -65 | 2 |
| Fast spiking(FS) | 0.1 | 0.2 | -65 | 2 |
| Bursting(BS) | 0.02 | 0.2 | -50 | 2 |

#### State-dependent Stochastic Bursting Neuron (SSBN)

For the Izhikevich neuron model as well as other similar models, the various possible firing patterns are tightly coupled to *f − I* curve of the neurons. Thus, the effects of firing patterns on network activity cannot be studied independently of the neuronal firing rate. To overcome this problem, we introduce a novel neuron model, the State-dependent Stochastic Bursting Neuron (SSBN). The SSB neuron has identical membrane potential dynamics as the Leaky Integrate and Fire (LIF) neuron given by

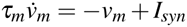

but the action-potential generation mechanism is stochastic. That is, whenever a predefined threshold *u_th_* is reached, *b* number of spikes are generated with probability 1/*b*. The inter-spike-interval within the burst is constant (2*ms*). The membrane potential is reset only after all spikes of the burst are produced. Thus, the SSBN neuron produces bursts of different lengths without altering the *f − I* curve. The simulation parameters are defined in **Table 2**.

**Table 2.**
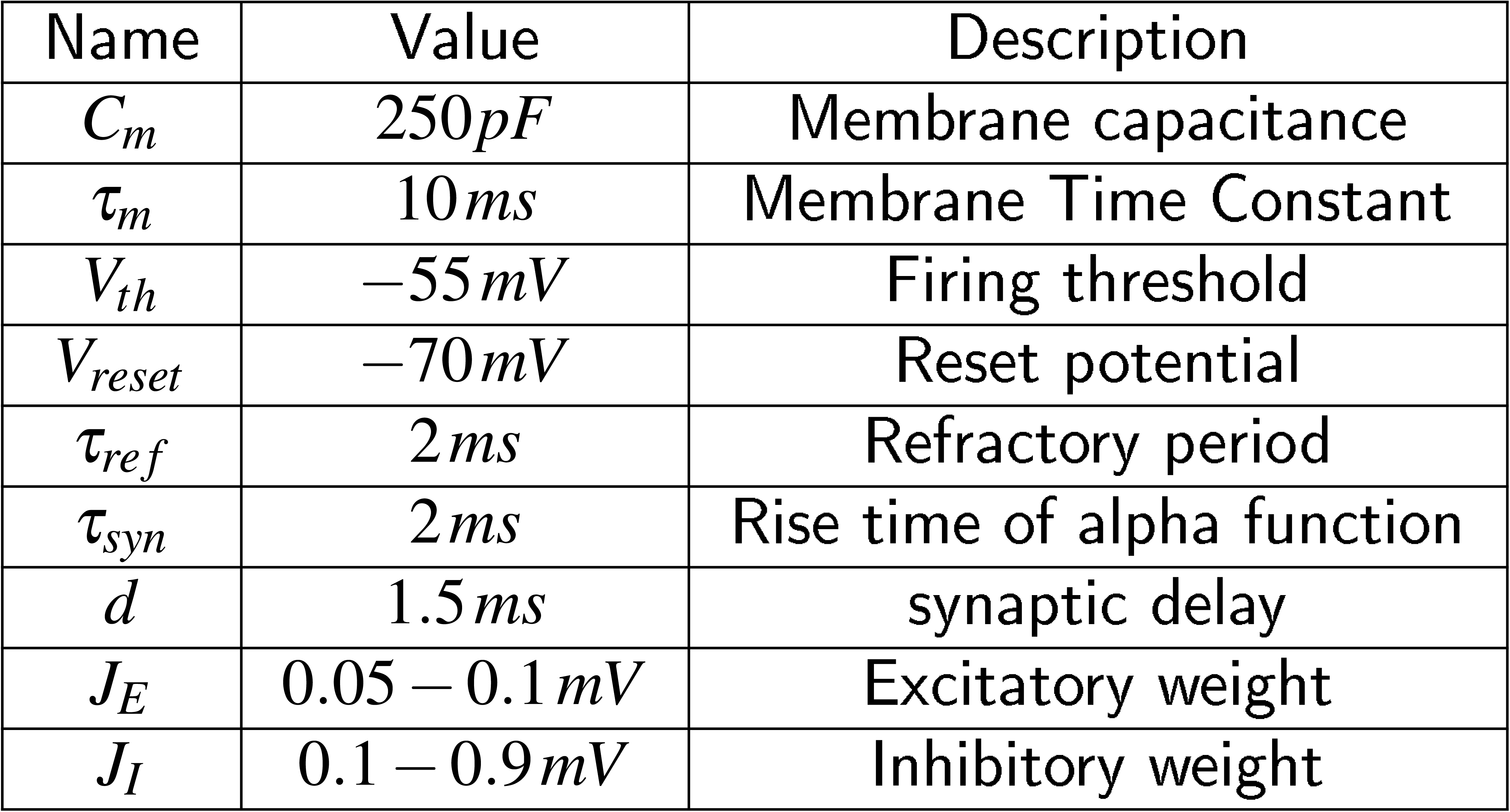
Simulation parameters

To make the above neuron model more biologically realistic, we let the number of spikes/burst *b* be a function of the mean input current that a neuron receives. The mean input current, *I_inp_* is a function of excitatory population firing rate, r i. e., 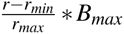, where *r_min_* is the firing rate of the population with minimum number of spikes per burst and *r_max_* is the population firing rate for the maximum number of spikes per burst in the inhibitory neurons, *B_max_.* More specifically, *b* is drawn from a binomial distribution (every 1000 ms) *b* ~ *B*(*n,p*) with mean *E*[*b*] *= f* (*I_inp_*) *= np, n* denotes the maximum number of spikes per burst which is fixed to *n* = 4 and *p* is the probability of producing one spike. Thus the mean input current to the neuron *I_inp_* affects the probability *p*. This we call the modified SSBN and this model is used in (Fig. 6 (inset)) only.

### Asynchronous state

In the stable asynchronous state the population activity is constant *r*(*t*) = *r_E_* = *r_I_* = *r*_0_. The mean recurrent input that each neuron receives is therefore also constant and given by

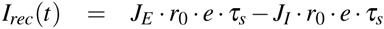

We study the stability of the asynchronous state following a linear perturbation approach^36,38^. A small oscillatory modulation of the stationary firing rate *r*(*t*) *= r*_0_ *+ r*_1_ *e^λt^* with *r*_1_ *≪* 1 and *λ = x* + *jω* where *ω* is the modulation frequency leads to corresponding oscillation of the synaptic current

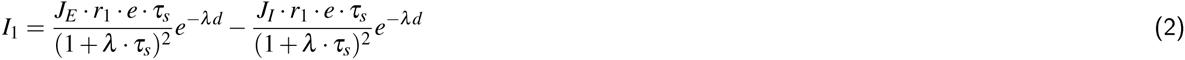

The firing rate in response to an oscillatory input is given by

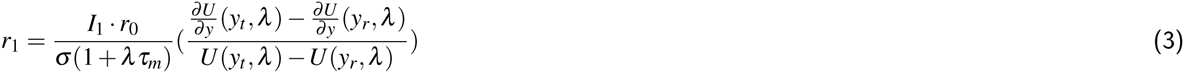

The function *U* is given in terms of combinations of hypergeometric functions

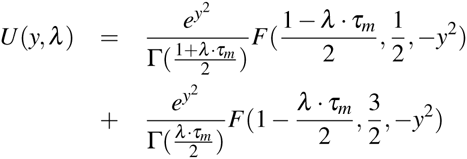

In a recurrent network the modulation of the firing rate and the modulation of the synaptic input must be consistent. Combining (2) and (3) we get

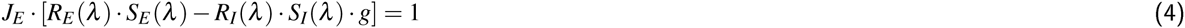

with

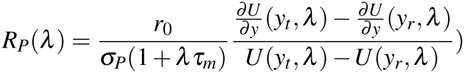

and

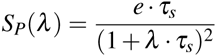

where *S_P_* is the synaptic response function for alpha-shaped postsynaptic currents

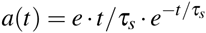

*P = E,I* denotes either the excitatory or inhibitory population.

If the inhibitory population is bursting the synaptic response function is given by

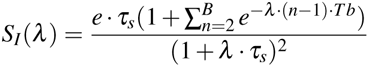

where *T_b_* is the length of the inter spike interval within a burst and *B* is the number of spikes in a burst. To compensate for the increased PSP due to bursting, the recurrent inhibitory firing rate is divided by *B*.

The critical coupling values at which modes have marginal stability with frequency *ω_i_* can then simply be computed by

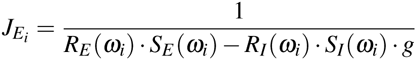

The smallest value 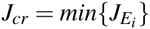 is the critical coupling value at which the first complex pair of eigenvalues crosses the imaginary axis and the system becomes unstable. The critical coupling values for different *B* is given by the dotted line in Fig. 5A.

### Networks

We generate networks of 4000 excitatory and 1000 inhibitory neurons randomly connected with a fixed probability of 0.1. In all simulations the excitatory neurons are of the regular-spiking type (*RS*), while the inhibitory neurons are divided into fast-spiking (*FS*) and bursting type (*BS*). The fraction of (*BS*) neurons is systematically varied between 0 and 1. For each network we compute the fraction of BS neurons, given by *F* = *N_bS_*/*N_i_*, with *N_I_ = N_FS_ + N_bS_*, where *N_FS_,N_bS_,N_I_* are the number of *FS, BS* and total number of inhibitory neurons respectively. Each neuron in the network receives poisson background input of rate *η*. The ratio of the synaptic strength of the excitatory and inhibitory connections is denoted by *g*.

### Hysteresis

To test the network response when network activity and spikes per burst were mutually dependent we changed the number of spikes per burst as a function of network firing rate. That is, at low firing rate, the network was composed only of non-bursting neurons. However, as the network output firing rate was increased by slowly increasing the external input was increased neurons started to burst. To implement a state-dependence of the burst size, we quantized the firing rate of the excitatory neurons into five non-overlapping ranges ([5 × (*B −* 1) − 5 × B] spikes/sec., where *B ϵ* {1,2,3,4,5}). The SSB neuron generated *B* spikes depending on the level of the excitatory firing rate. To change the number of spikes per burst, we estimated the input rate either in 3sec (Fig. 6A) or 200ms windows (Fig. 6B). To change the network firing rate, we changed the external input to the network in steps of 100spikes/sec every 3sec (Fig. 6A) or 200 ms (Fig. 6B). The external input was varied until the BS neurons reached a maximal burst size *B* = 5), after that the external input was reduced with the same rate.

### Data Analysis

We use the mean firing rate (*v*) and Fano facor (*FF*) to characterise the dynamical states of the networks. Mean firing rate is measured as the number of spikes per neuron per second. *FF* is used to quantify the synchrony in the network. The *FF* of a population is defined as

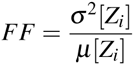

To obtain a reliable estimate of the population activity, the cumulative activity of the spike trains of all the neurons in the network were binned in discrete time bins(bin width = 2ms). *Z_i_* is the population activity in a bin *i*. An increase in positive correlation increased the *Variance*[*Z_i_*] and consequently the *FF*[*Z_i_*].

Coefficient of variation, *CV*, of the inter-spike interval distribution T of a neuron, is given by

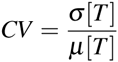

The mean *CV* of the neurons in a population gives the regularity of neuronal spiking in the population.

#### Spectral Entropy

To quantify the degree of oscillatory activity in a network we compute the spectral entropy *H_S_*, which is a measure of dispersion of spectral energy of a signal.^54^ It is given by

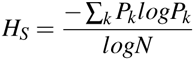

where *P_k_* is the spectral power at frequency *k* and *N* is the total number of frequency bins considered. The power spectrum is computed using a Fast-Fourier-Transform of the population activity *v* and normalized such that *Σ_k_ p_k_* = 1.

A flat power spectrum, e.g. in the white noise case, has maximum spectral entropy, i.e. *H_S_* = 1. By contrast, a spectrum with all power concentrated in one frequency, e.g. periodic sine signal, has zero spectral entropy *H_S_* = 0. Therefore, the more oscillatory the activity dynamics is, the smaller *H_S_* will be. In our simulations, the value of spectral entropy ranged from 0.2 to 0.9.

While Fano factor is a good descriptor of the synchronicity in the network activity, it does not quantify network oscillations. Whenever, we wanted to quantify the strength of the network oscillations specifically, we have used spectral entropy.

### Simulation and Data Analysis Tools

All network simulations are written in Python (http://www.python.org) and implemented in NEST (http://www.nest-initiative.org).^55^ A temporal resolution of 0.1 ms is used for the integration of the differential equations. Results were analyzed using SciPy and NumPy libraries. Visualizations were done using Matplotlib.^56^

## Acknowledgements

We thank Dr.Christopher Kim for comments on an earlier version of this manuscript. This work was supported by the German Federal Ministry of Education and Research (01GQ0830 to BFNT Freiburg/Tuebingen), FACETS-ITN (PITN-GA-2009-237955) and the BrainLinks-BrainTools Cluster of Excellence funded by the German Research Foundation (DFG #EXC 1086). AK and AS acknowledge INTERREG IV Rhin superieur program and European Funds for Regional Development (FEDER) through the project TIGER A31. We also acknowledge the use of the computing resources provided by the Black Forest Grid Initiative and the bwGRiD (http://www.bw-grid.de), member of the German D-Grid initiative, funded by the Ministry for Education and Research and the Ministry for Science, Research and Arts Baden-Wuerttemberg.

## Additional Information

The project was funded in part by following funding agencies: Bundesministerium fuer Forschung und Technologie (German Ministry for Research and Technology) - 01GQ0830 [to Ad Aertsen and Arvind Kumar]; Deutsche Forschungsgemeinschaft (DFG) - EXC 1086 [to Ad Aertsen and Arvind Kumar]; Ajith Sahasranamam was partially funded by the FACETS-ITN (PITN-GA-2009-237955). AK and AA also acknowledge the INTERREG IV Rhin Superieur Program and the European Funds for Regional Development (FEDER) through the project TIGER A31. We also acknowledge the use of the computing resources provided by the Black Forest Grid Initiative and the bwGRiD (http://www.bw-grid.de), member of the German D-Grid initiative, funded by the Ministry for Education and Research and the Ministry for Science, Research and Arts Baden-Wuerttemberg.

## Author Contributions

Conceived and designed the experiments: AS, IV, AA, AK. Performed the experiments: AS, IV. Analyzed the data: AS, IV. Contributed reagents/ materials/analysis tools: AK. Contributed to the writing of the manuscript: AS, IV, AA, AK.

## Supplementary figures

**Figure S1.** Increased bursting reduces the frequency of input oscillations that can be tracked. In unconnected networks of different number of modified-SSBNs, we test how well sinusoidally modulated input of different frequencies could be followed by the population. For higher input frequencies, it is seen that for increased number of spikes per burst are less able to follow the input. The rasters and the z-scored PSTHs for different number of spikes per burst for a fixed sinusoidal input (120 Hz) are shown in (**A**) and (**B**). (**C**) For a fixed size of the neuron ensemble (*N* = 100) it is observed that the normalized power of the peak frequency drops and saturates at a very low value (≈ 0) for higher frequencies of the sinusoidally modulated input. (**D**) The map shows the maximum frequency of the input that can be tracked by different combinations of number of independent neurons in the population and the number of spikes per burst. While the value of the frequency drops with the increase in the number of spikes per burst, it can be compensated for by increasing the number of neurons in the ensemble.

**Figure S2.** Effect of addition of bursting neurons on the state of network composed of SSBN. (A) Evolution of spectral entropy(*H_S_*) for a network which is initially synchronous and changes to being asynchronous with the addition of bursting neurons, added

(g = 11,*d* = 2*ms, η* = 10500 – 11500 *sp/s, J_e_* = 0.1 mV).(**B**) a network in an asynchronous state that continues to remain asynchronous with the addition of bursting neurons

(g = 5, *d* = 2.0*ms, η* = 4000 – 5000*sp/s, J_E_* = 0.04*mV*), **C** qualitatively synchronous activity can remain synchronous even when inhibitory neuron firing patterns are changed

(g = 8, *d* = 4*ms, η* = 8500 – 9500*sp/s, J_E_* = 0.1 *mV*) and (**D**) an initially asynchronous activity in the network that becomes synchronous with the addition of bursting neurons

(g = 6, *d* = 2 *ms, η* = 4500 – 5500*sp/s, J_E_* = 0.1 *mV*). (**E**) The Fano Factor values of the different transitions are plotted against the changes in the fraction of bursting neurons. The different colours correspond to the different state transitions observed (colours marked in the titles of **A,B,C** and D.)(F) The rasters illustrating the four types of transitions are shown in a phase space of *FF* and the difference in *H_s_.* The difference in *H_s_* is the difference in spectral entropy between the initial and final points of each transitions. The initial rasters are marked in yellow and the final rasters are marked in black in the corresponding panels **A,B,C** and **D**. The *FF* values marked are the *FF* values of the initial points.

**Figure S3.** A simple network producing an external input induced spiking of a presynaptic BS Population. This BS population acted as the inhibitory presynaptic input to a regular LIF neuron. The membrane potential of this LIF neuron was maintained very close to the threshold by an external poissonian input. The percentage change in the variance of the membrane potential (**A**) and firing rate (**B**) of the postsynaptic LIF neuron with the varying number of spikes per burst in the presynaptic SSBN population is plotted. The increase in the size of the presynaptic population decreased the amount of changes in the variance of the membrane potential and the firing rate of the post-synaptic LIF with the change in the number of spikes per burst.

**Figure S4.** Burstiness of single neurons changes with network state. The number of spikes per burst that a BS neuron(lzhikevich model) produces depends on the state of the network. To quantify the burstiness of a neuron we use the Bursting Index.^57^ This measure assigns a rank *R_n_* to every interspike interval (ISI) of a spike train. The lowest value of an ISI has zero rank. If the ISIs are independent, the value of each ISI can be considered to be a random number drawn from a uniform distribution between 1 and N, where N is the total number of ISIs. If a spike train contains a burst, then this assumption does not hold anymore. The Bursting Index is equivalent to the Rank Surprise (RS) statistic, which captures the discrepancy between the case of having independent and uniformly distributed sequence of variables *R_n_,…,R_n+q−1_* and the actual outcome in the case of a burst consisting of *q* number of spikes. It is given given by *RS = −log*(*P*(*T_q_ ≤ r_n_* +…+ *r_n+q−1_*)) *where r_n_* is the observed value of rank *R_n_. T_q_* is the sum of *q* discrete uniform variates between 1 and N. In the above figure, the average bursting index of BS neurons for different *η* and *g* values are shown in a randomly connected network of excitatory-BS neurons (Izhikevich model)

## References

1. Markram, H. et al. Interneurons of the neocortical inhibitory system. Nat Rev Neurosci 5, 793–807 (2004).

2. Luo, L., Callaway, E. M. & Svoboda, K. Genetic dissection of neural circuits. Neuron 57, 634–60 (2008).

3. Defelipe, J. et al. New insights into the classification and nomenclature of cortical gabaergic interneurons. Nat Rev Neurosci 14, 202–216 (2013).

4. Wichterle, H., Gifford, D. & Mazzoni, E. Mapping neuronal diversity one cell at a time. Science 341, 726–727 (2013).

5. Neske, G. T., Patrick, S. L. & Connors, B. W. Contributions of diverse excitatory and inhibitory neurons to recurrent network activity in cerebral cortex. J Neurosci. 35, 1089–1105 (2015).

6. Yizhar, O. et al. Neocortical excitation/inhibition balance in information processing and social dysfunction. Nature 477, 171–8 (2011).

7. Sohal, V. S., Zhang, F., Yizhar, O. & Deisseroth, K. Parvalbumin neurons and gamma rhythms enhance cortical circuit performance. Nature 459, 698–702 (2009).

8. Cardin, J. A. et al. Driving fast-spiking cells induces gamma rhythm and controls sensory responses. Nature 459, 663–7 (2009).

9. Wilson, N. R., Runyan, C. A., Wang, F. L. & Sur, M. Division and subtraction by distinct cortical inhibitory networks in vivo. Nature 488, 343–348 (2012).

10. Denker, M., Timme, M., Diesmann, M., Wolf, F. & Geisel, T. Breaking synchrony by heterogeneity in complex networks. Phys Rev Lett 92, 074103–1–074103-4 (2004).

11. Padmanabhan, K. & Urban, N. N. Intrinsic biophysical diversity decorrelates neuronal firing while increasing information content. Nat Neurosci 13, 1276–82 (2010).

12. Pinto, L. & Dan, Y. Cell-Type-Specific Activity in Prefrontal Cortex during Goal-Directed Behavior. Neuron 87, 437–450 (2015).

13. Diester, I. et al. An optogenetic toolbox designed for primates. Nat Neurosci 14, 387–97 (2011).

14. Achard, P. & Schutter, E. D. Complex parameter landscape for a complex neuron model. PLoS Comput Biol 2, e94 (2006).

15. Prinz, A. A., Bucher, D. & Marder, E. Similar network activity from disparate circuit parameters. Nat Neurosci 7, 1345–1352 (2004).

16. Marder, E. & Taylor, A. L. Multiple models to capture the variability in biological neurons and networks. Nature 14, 133–138 (2011).

17. Ascoli, G. A. et al. Petilla terminology: nomenclature of features of GABAergic interneurons of the cerebral cortex. Nature Reviews Neuroscience 9, 557–568 (2008).

18. Gupta, A., Wang, Y. & Markram, H. Organizing principles for a diversity of GABAergic interneurons and synapses in the neocortex. Science 287, 273–278 (2000).

19. Jarsky, T., Mady, R., Kennedy, B. & Spruston, N. Distribution of bursting neurons in the CA1 region and the subiculum of the rat hippocampus. J Comp Neurol 506, 535–547 (2008).

20. Larkum, Μ. E., Zhu, J. J. & Sakmann, B. Dendritic mechanisms underlying the coupling of the dendritic with thr axonal action potential initiation zone of adult rat layer 5 pyramidal neurons. J Physiol 533, 477–466 (2001).

21. Womelsdorf, T., Ardid, S., Everling, S. & Valiante, T. A. Burst firing synchronizes prefrontal and anterior cingulate cortex during attentional control. Current Biology 1–9 (2014).

22. Sanabria, E. R., Su, H. & Yaari, Y. Initiation of network bursts by Ca2+-dependent intrinsic bursting in the rat pilocarpine model of temporal lobe epilepsy. J Physiol (London) 532, 205–216 (2001).

23. Tachibana, Y., Iwamuro, H., Kita, H., Takada, M. & Nambu, A. Subthalamo-pallidal interactions underlying parkinsonian neuronal oscillations in the primate basal ganglia. Eur J Neurosci 34, 1470–1484 (2011).

24. Markram, H., Wang, Y. & Tsodyks, M. Differential signaling via the same axon of neocortical pyramidal neurons. Proc Natl Acad Sci USA 95, 5323–5328 (1998).

25. Wittenberg, G. M. & Wang, S. S.-H. Malleability of spike-timing-dependent plasticity at the ca3-ca1 synapse. J Neurosci 26, 6610–6617 (2006).

26. Kumar, A. & Mehta, M. R. Frequency dependent changes in nmdar-dependent synaptic plasticity. Front Comput. Neurosci 5 (2011).

27. Wang, X.-J. Neurophysiological and computational principles of cortical rhythms in cognition. Physiological Reviews 90, 1195–268 (2010).

28. Bogaard, A., Parent, J., Zochowski, M. & Booth, V. Interaction of cellular and network mechanisms in spatiotemporal pattern formation in neuronal networks. J Neurosci 29, 1677–1687 (2009).

29. Krahe, R. & Gabbiani, F. Burst firing in sensory systems. Nat. Rev. Neurosci. 5, 13–23 (2004).

30. Izhikevich, E. M. Simple mode of spiking neurons. IEEE Tran, on Neural Networks 14, 1569–1572 (2003).

31. Uhlhass, P. J. et al. Neural synchrony in cortical networks: history, concept and current status. Front Integr Neurosci 3, 1–19 (2009).

32. Fries, P. A mechanism for cognitive dynamics: neuronal communication through neuronal coherence. Trends in Cognitive Sciences 9, 474–480 (2005).

33. Buzsáki, G. & Wang, X.-J. Mechanisms of gamma oscillations. Annu Rev Neurosci 35, 203–225 (2012).

34. Brunei, N. & Wang, X.-J. What determines the frequency of fast network oscillations with irregular neural discharges? i. synaptic dynamics and excitation-inhibition balance. J Neurophysiol 90, 415–430 (2003).

35. Ledoux, E. & Brunel, N. Dynamics of networks of excitatory and inhibitory neurons in response to time-dependent inputs. Front Comput Neurosci 5, 1–17 (2011).

36. Brunei, N. & Hakim, V. Sparsely synchronized neuronal oscillations. Chaos 18, 015113 (2008).

37. Tiesinga, P. & Sejnowski, T. J. Cortical enlightenment: Are attentional gamma oscillations driven by ing or ping? Neuron 63, 727–732 (2009).

38. Brunei, N. & Hansel, D. How noise affects the synchronization properties of recurrent networks of inhibitory neurons. Neural Computation 18, 1066–110 (2006).

39. Brunei, N. Dynamics of sparsely connected networks of excitatory and inhibitory spiking neurons. J Comput Neurosci 8, 183–208 (2000).

40. Kumar, A., Schrader, S., Aertsen, A. & Rotter, S. The high-conductance state of cortical networks. Neural Computation 20, 1–43 (2008).

41. Hahn, G., Bujan, A. F., Frégnac, Y., Aertsen, A. & Kumar, A. Communication through resonance in spiking neuronal networks. PloS Comput Biol 1–16 (2014).

42. vanVreeswijk, C. & Sompolinsky, H. Chaos in neuronal networks with balanced excitatory and inhibitory activity. Science 274, 1724–1726 (1996).

43. Stern, M., Sompolinsky, H. & Abbott, L. F. Dynamics of random neural networks with bistable units. Phys. Rev. E 90, 1–7 (2014).

44. Amit, D. & Brunel, N. Model of global spontaneous activity and local structured activity during delay periods in the cerebral cortex. Cereb. cortex 7, 237–252 (1997).

45. Mongillo, G., Hansel, D. & van Vreeswijk, C. Bistability and spatiotemporal irregularity in neuronal networks with nonlinear synaptic transmission. Phys. Rev. Lett. 108, 158101 (2012).

46. Schnepel, P., Kumar, A., Zohar, M., Aertsen, A. & Boucsein, C. Physiology and impact of horizontal connections in rat neocortex. Cerebral Cortex 1–18 (2014).

47. Bonifazi, P. et al. Gabaergic hub neurons orchestrate synchrony in developing hippocampal networks. Science 326, 1419–1424 (2009).

48. Vlachos, I., Aertsen, A. & Kumar, A. Beyond statistical significance: Implications of network structure on neuronal activity. PLoS Comput Biol 8, e1002311 (2012).

49. Kumar, A., Vlachos, I., Aertsen, A. & Boucsein, C. Challenges of understanding brain function by selective modulation of neuronal subpopulations. Trends in Neurosciences 36, 579–586 (2013).

50. Gutierrez, G. J. & Marder, E. Rectifying electrical synapses can affect the influence of synaptic modulation on output pattern robustness. J Neurosci 33, 13238–13248 (2013).

51. Connors, B. W. & Gutnick, M. J. Intrinsic firing patterns of diverse neocortical neurons. Trends in Neurosciences 13, 99–104 (1990).

52. Karimi, F. & Holme, P. Threshold model of cascades in empirical temporal networks. Physica A: Statistical Mechanics and its Applications 392, 3476–3483 (2013).

53. Vogels, T. P., Sprekeler, H., Zenke, F., Clopath, C. & Gerstner, W. Inhibitory plasticity balances excitation and inhibition in sensory pathways and memory networks. Science 334, 1569–1573 (2011).

54. Blanco, S., Garay, A. & Coulombie, D. Comparison of frequency bands using spectral entropy for epileptic seizure prediction. ISRN Neurol 2013, 287327 (2013).

55. Gewaltig, M.-O. & Diesmann, M. Nest (neural simulation tool). Scholarpedia 2, 1430 (2007).

56. Hunter, J. D. Matplotlib: A 2d graphics environment. Computing In Science & Engineering 9, 90–95 (2007).

57. Gourévitch, B. & Eggermont, J. J. A nonparametric approach for detection of bursts in spike trains. J Neurosci Methods 160, 349–58 (2007).

